# Screening and directed evolution of transporters to improve xylodextrin utilization in the yeast *Saccharomyces cerevisiae*

**DOI:** 10.1101/166678

**Authors:** Chenlu Zhang, Ligia Acosta-Sampson, Vivian Yaci Yu, Jamie H. D. Cate

## Abstract

The economic production of cellulosic biofuel requires efficient and full utilization of all abundant carbohydrates naturally released from plant biomass by enzyme cocktails. Recently, we reconstituted the *Neurospora crassa* xylodextrin transport and consumption system in *Saccharomyces cerevisiae*, enabling growth of yeast on xylodextrins aerobically. However, the consumption rate of xylodextrin requires improvement for industrial applications, including consumption in anaerobic conditions. As a first step in this improvement, we report analysis of orthologues of the *N. crassa* transporters CDT-1 and CDT-2. Transporter ST16 from *Trichoderma virens* enables faster aerobic growth of *S. cerevisiae* on xylodextrins compared to CDT-2. ST16 is a xylodextrin-specific transporter, and the xylobiose transport activity of ST16 is not inhibited by cellobiose. Other transporters identified in the screen also enable growth on xylodextrins including xylotriose. Taken together, these results indicate that multiple transporters might prove useful to improve xylodextrin utilization in *S. cerevisiae*. Efforts to use directed evolution to improve ST16 from a chromosomally-integrated copy were not successful, due to background growth of yeast on other carbon sources present in the selection medium. Future experiments will require increasing the baseline growth rate of the yeast population on xylodextrins, to ensure that the selective pressure exerted on xylodextrin transport can lead to isolation of improved xylodextrin transporters.

## Introduction

Cellulosic biofuel production from plant biomass requires efficient use of all abundant carbohydrates in the plant cell wall [1, 2]. To make the fermentation process economically feasible, engineered yeast should be able to consume a mixture of sugars naturally released by enzyme cocktails simultaneously, including hexose and pentose sugars [3-5]. Xylodextrins (XDs), such as xylobiose and xylotriose, are oligomers of β-1,4-linked xylose. They are derived from hemicellulose, one of the major forms of biomass in lignocellulose, and are hydrolyzed to xylose by β-xylosidase. The first example of ethanol production from xylan was reported in 2004, wherein xylanase and β-xylosidase were displayed on the cell surfaces of *S. cerevisiae* expressing a xylose consumption pathway [6]. However, the xylan degrading and xylose utilization abilities of this recombinant *S. cerevisiae* strain requires further optimization to be industrially useful. Recently, a XD utilization pathway from *N. crassa*, which requires a transporter CDT-2 along with two intracellular β-xylosidases GH43-2 and GH43-7, was identified and subsequently engineered into *S. cerevisiae*, enabling the yeast to grow aerobically on XD, or co-ferment XDs with xylose or other hexose sugars [7]. However, the consumption rate of XDs remains quite slow and thus needs to be improved for future industrial applications.

The transporter CDT-2 belongs to the Major Facilitator Superfamily (MFS), one of the largest and most ubiquitous secondary transporter subfamilies [8]. Its members exist in all species from bacteria to mammals, and they range in size from 400 to 600 amino acids, organized into 12 trans-membrane α-helices [9]. CDT-2 belongs to the hexose family of MFS transporters, which includes yeast hexose transporters, human glucose transporters, and xylose transporters [10-13].

For industrial applications, naturally occurring transporters are generally not optimal without further engineering. Protein engineering, including rational design and directed evolution approaches, has been widely used to improve the performance of a wide variety of enzymes and pathways. However, engineering membrane proteins such as MFS transporters has proven to be challenging [14-16]. Directed evolution is an important means for carrying out protein engineering [17], but requires developing a high-throughout screening system [18, 19]. In 2014, Ryan et al. developed a new method for performing directed evolution experiments in yeast using the CRISPR-Cas9 system [20]. Integrating linear DNA into the genome by homologous recombination mediated CRISPR-Cas9 overcame high levels of copy number variation seen with plasmid-based expression, and allowed directed evolution of CDT-1 to isolate an improved cellodextrin transporter [20].

To improve xylodextrin utilization in *S. cerevisiae*, we sought to identify potential XD transporters by characterizing a library of CDT-1 and CDT-2 orthologues. We analyzed cellular localization, transport activity and aerobic growth profiles. Moreover, with one of the best performing transporters identified in the screen, we attempted to use CRISPR-Cas9 mediated directed evolution to improve XD consumption in yeast.

## Results

To identify XD transporters that could be used to improve utilization of xylodextrins, we carried out a screening of codon-optimized CDT-1 and CDT-2 transporter orthologues, named STX (X spanning from 1 to 17) (S1 Table). Phylogenetic analysis suggested that orthologues of CDT-2 are widely distributed in the fungal kingdom, indicating that many fungi are able to consume xylodextrins derived from plant cell walls (Figure 1). These orthologues are 28-67% identical in amino acid sequence to CDT-2.

**Figure 1.**
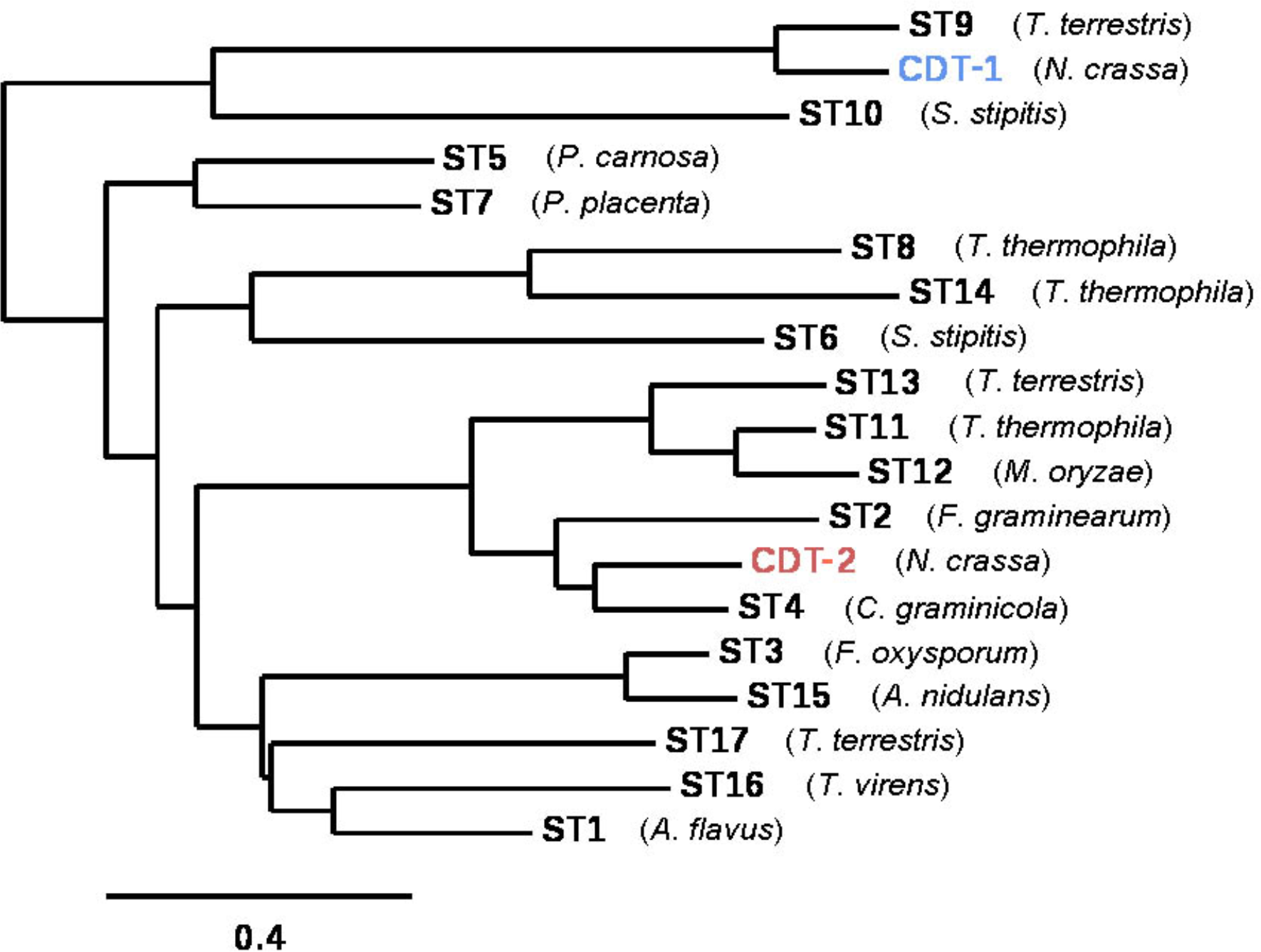
Phylogenetic tree of CDT-1 and CDT-2 related transporters. The phylogenetic tree was constructed using the Phylogeny.fr platform and the amino acid sequences for the transporters used in this study. [http://www.phylogeny.fr/index.cgi, [31, 32]]. The species name from which each transporter was identified is indicated, and protein sequences are included in (S1 Table).

We first constructed plasmids expressing ST1-ST17 fused to enhanced GFP at the C-terminus, using the strong *TDH3* promoter and the *CYC1* terminator, and expressed the transporters in *S. cerevisiae* strain D452-2. Using epi-fluorescent microscopy, we found that most transporters localize to the plasma membrane (Figure 2). Transporters ST5, ST8, ST10 and ST14 are dispersed in the cytoplasm or vacuole (Figure 2), indicating these transporters are not well folded in *S. cerevisiae*, and they were not analyzed further.

**Figure 2.**
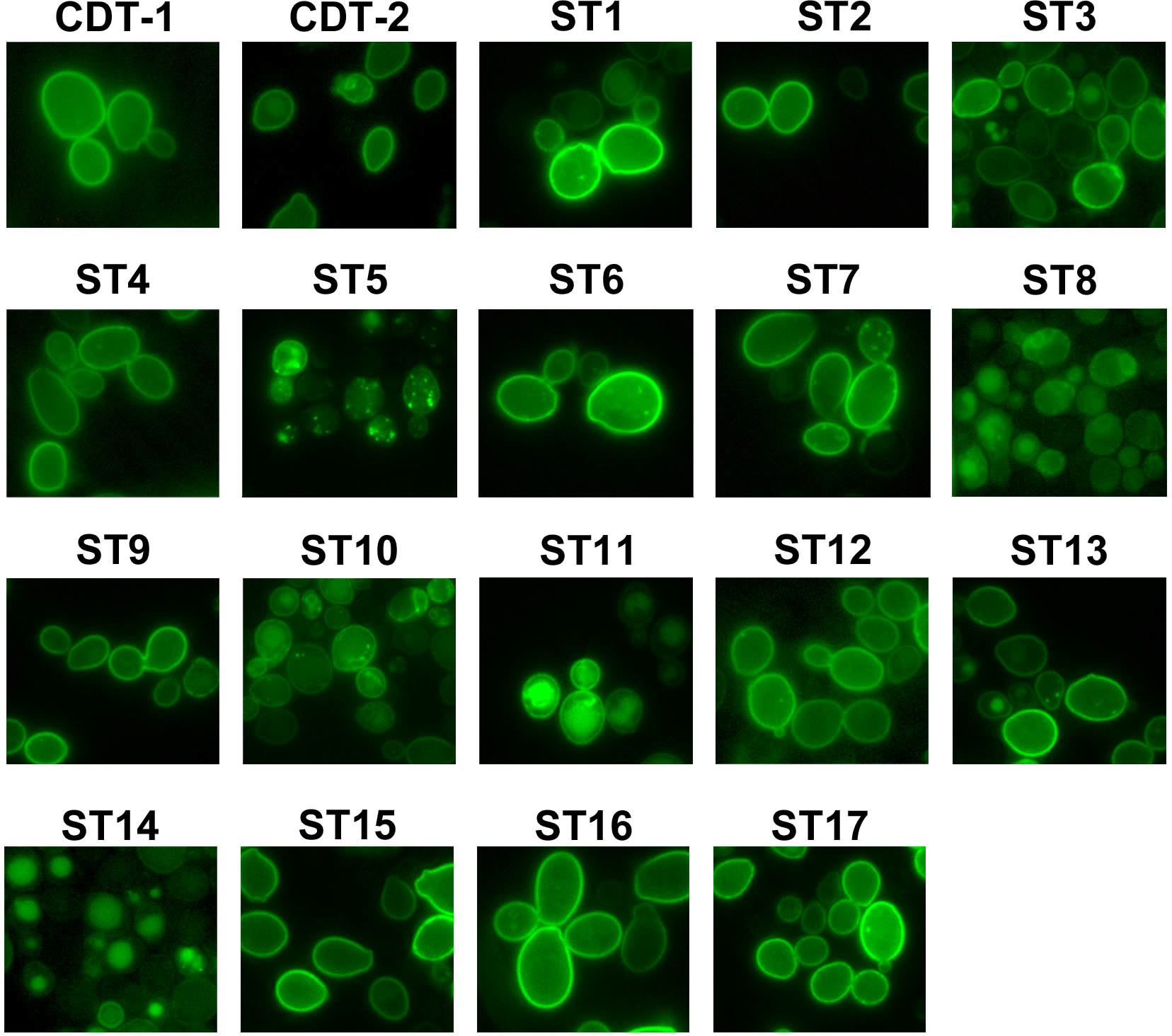
Cellular localization of transporters. Images of *S. cerevisiae* strain D452-2 cells with CDT-1–GFP, CDT-2–GFP and STX–GFP (ST1 to ST17) transporters using epi-fluorescent microscopy. Transporters mainly localize to the plasma membrane, except for ST5, ST8, ST10 and ST14.

Xylodextrins, including xylobiose and xylotriose, are β-1,4-linked xylose oligomers with different degree of polymerization. As a direct test of transport activity for cellobiose (G2), xylobiose (X2) and xylotriose (X3), we chose transporters that localized to the plasma membrane (Figure 2) and conducted a yeast cell-based sugar uptake assay. Notably, we found that ST3, ST15 and ST16 are XD-specific transporters, while other transporters showing XD transport activity are able to transport cellobiose as well, although the overall activity varied (Figure 3). The uptake activity of CDT-2 for both X2 and X3 was more than ten times that of ST9, ST11 and ST12 (Figure 3). Thus, these transporters were not studied further. The sugar preferences for ST transporters are summarized and shown on the phylogenetic tree in Figure S1.

**Figure 3.**
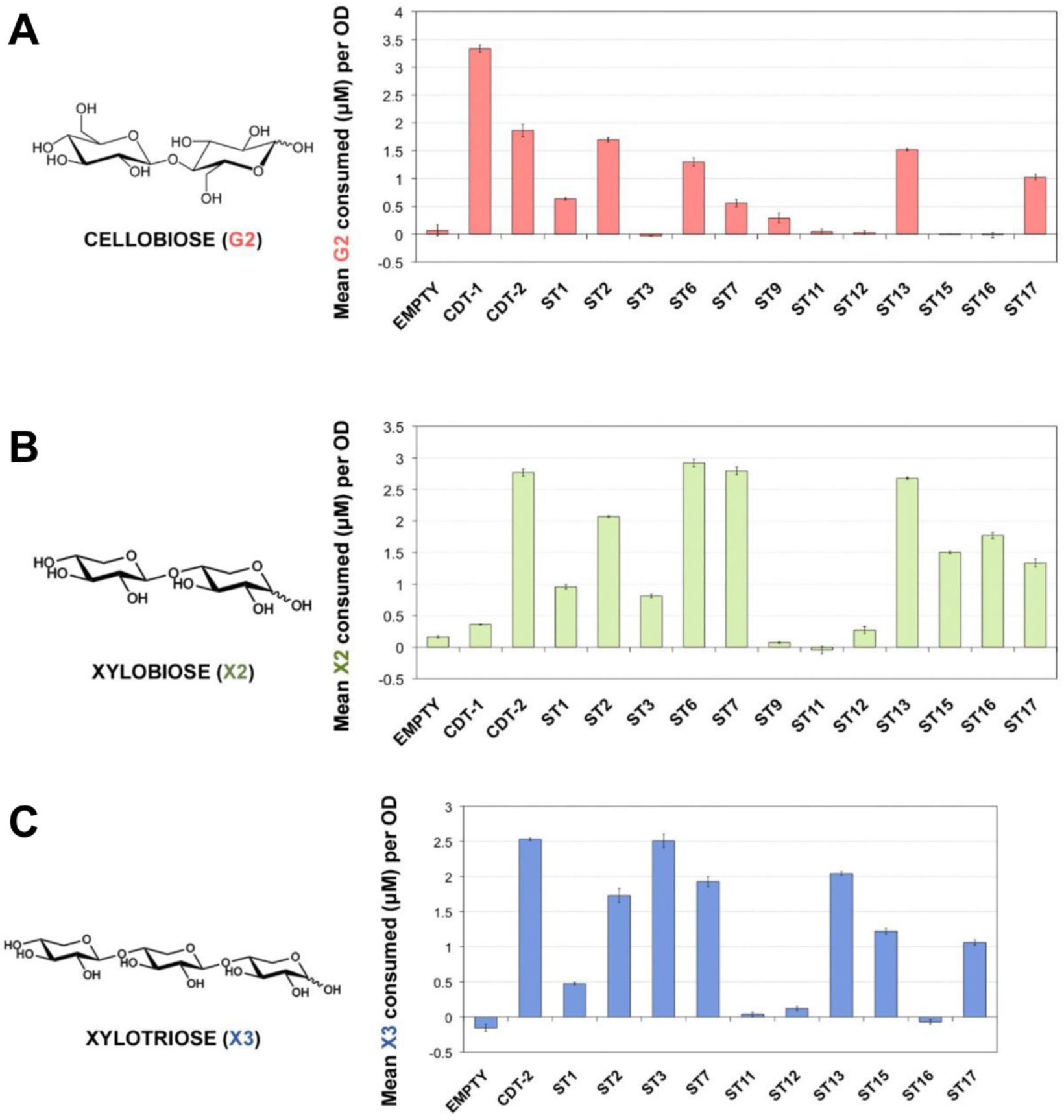
Cellobiose and xylobiose uptake of ST transporters *in vivo*. **A**) Cellobiose (G2), **B**) xylobiose (X2) and **C**) xylotriose (X3) uptake from D452-2 cells expressing ST transporters. Cells were incubated in buffer containing 200 µM sugar. Polysaccharide concentrations were measured at t = 0 min and t = 30 min. The concentration of sugar consumed was normalized to 1 OD unit of cells. Measurements represent the mean sugar uptake along with corresponding standard error from biological triplicates.

Many transporters are inhibited by the presence of related sugars in the extracellular medium. For example, xylose transporters are inhibited by the presence of glucose, which has led to many efforts to relieve this inhibition [21, 22]. We had previously observed that cellobiose inhibits xylobiose transport by CDT-2 [7]. We therefore tested xylobiose transport activity in the presence of increasing molar concentrations of cellobiose, up to 10 times the concentration of X2. Interestingly, we found that the uptake of xylobiose by ST15 and ST16 was not inhibited by cellobiose (Figure 4), making them good candidates for co-fermentation with glucose or cellodextrin from lignocellulosic biomass under a number of pretreatment scenarios. However, for other transporters, there was less X2 consumed per OD when increasing molar concentration of G2 were present in the media, i.e. as observed for CDT-2 (Figure 4).

**Figure 4.**
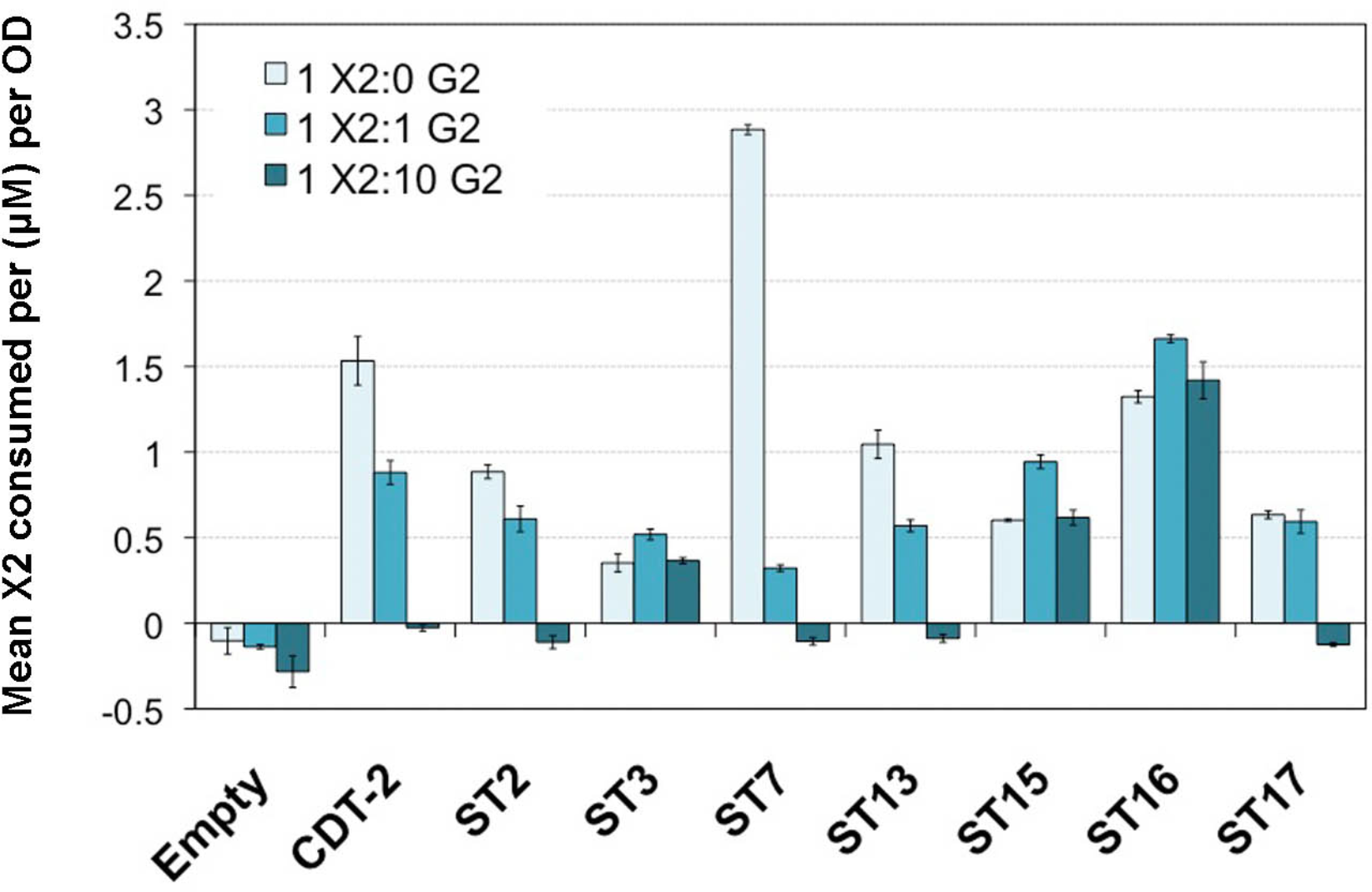
Competition between xylobiose and cellobiose uptake. Xylobiose uptake in the presence of increasing concentrations of cellobiose in cells expressing ST transporters. Cells were incubated in buffer containing 200 μM X2 along with increasing stoichiometric ratios of G2: 0X, 1X, and 10X the concentration of X2. Measurements represent mean X2 consumed after 30 min incubation from biological triplicates for each ratio indicated, along with standard error bars.

To further correlate the sugar uptake activity with yeast growth, we engineered the xylose-consuming *S. cerevisiae* strain SR8 [23] using CRISPRm (S2 Figure), by insertion of GH43-2 and GH43-7 at the *TRP1* and *LYP1* loci, respectively. The engineered SR8 strain was named SR8A. Yeast strain SR8A cells with plasmids expressing different transporters were cultured in oMM media under aerobic conditions for 96 hours using xylodextrin as the only carbon source. ST2 and ST16 enabled *S. cerevisiae* to grow more rapidly on xylodextrin (Figure 5). The area under the curve (AUC) of ST2 and ST16 were 1.16 and 1.26 times higher than CDT-2, respectively. We also tested the anaerobic growth profile of ST2 and ST16 transporter on XD plus sucrose, but there were no significant improvements compared with CDT-2 in anaerobic conditions.

**Figure 5.**
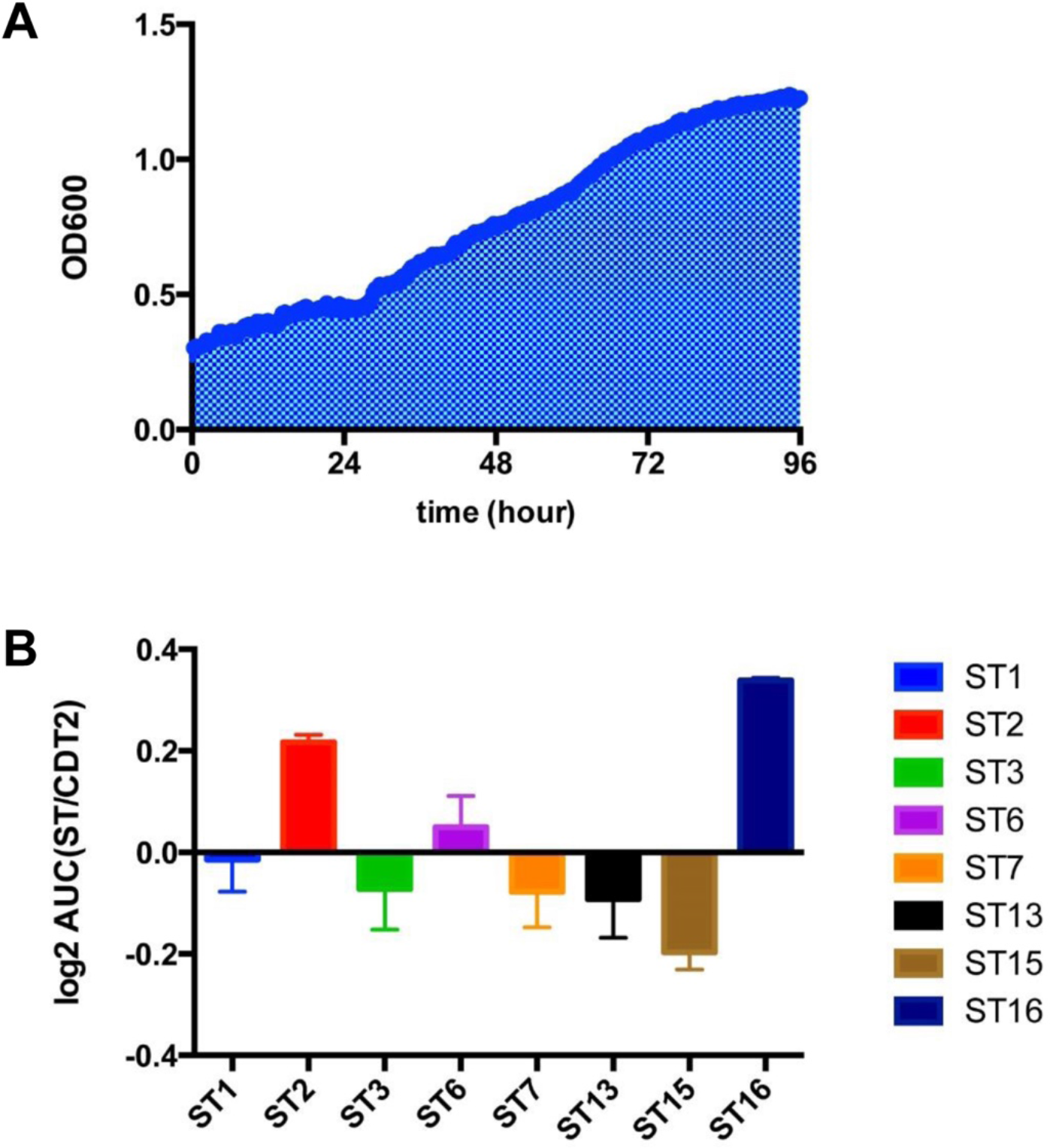
Aerobic growth profile of ST transporters on xylodextrin. **A**). Growth curve of SR8A strain expressing CDT-2 from a plasmid up to 96 hours, with the area under the curve (AUC) shaded. A representative experiment is shown with xylodextrin as the sole carbon source under aerobic conditions. **B**). AUCs for SR8A strains with plasmids expressing different ST transporters. AUCs are plotted as the log_2_ of the ratio of ST over CDT-2 AUCs. All experiments were conducted in biological triplicate, with error bars representing standard deviations.

Taken together, three potentially useful xylodextrin transporters were identified from the screen, each with different advantages and disadvantages (S1 Table). Of these, ST16 is a XD-specific transporter, showing faster aerobic growth and its xylobiose transport activity is not inhibited by cellobiose. We therefore chose ST16, which is derived from *Trichoderma virens,* as the starting point for setting up a directed evolution experiment to improve XD consumption. Based on experiments published previously, in which XD consumption was rapidly turned on and off in anaerobic conditions [7], we hypothesized that XD transport is limiting in terms of consumption rate. Notably, when we inserted the transporter ST16 under control of *PGK1* promoter and *TPS3* terminator into the yeast genome at the *LEU2* locus, there was a dramatic decrease in the growth rate compared with overexpression ST16 from a plasmid. Furthermore, the strain with incorporated ST16 grew almost as slow as the control (Figure 6A). GFP fluorescence emission spectroscopy verified that ST16 was expressed from the chromosome, but at lower levels than overexpression from a plasmid (Figure 6B).

**Figure 6.**
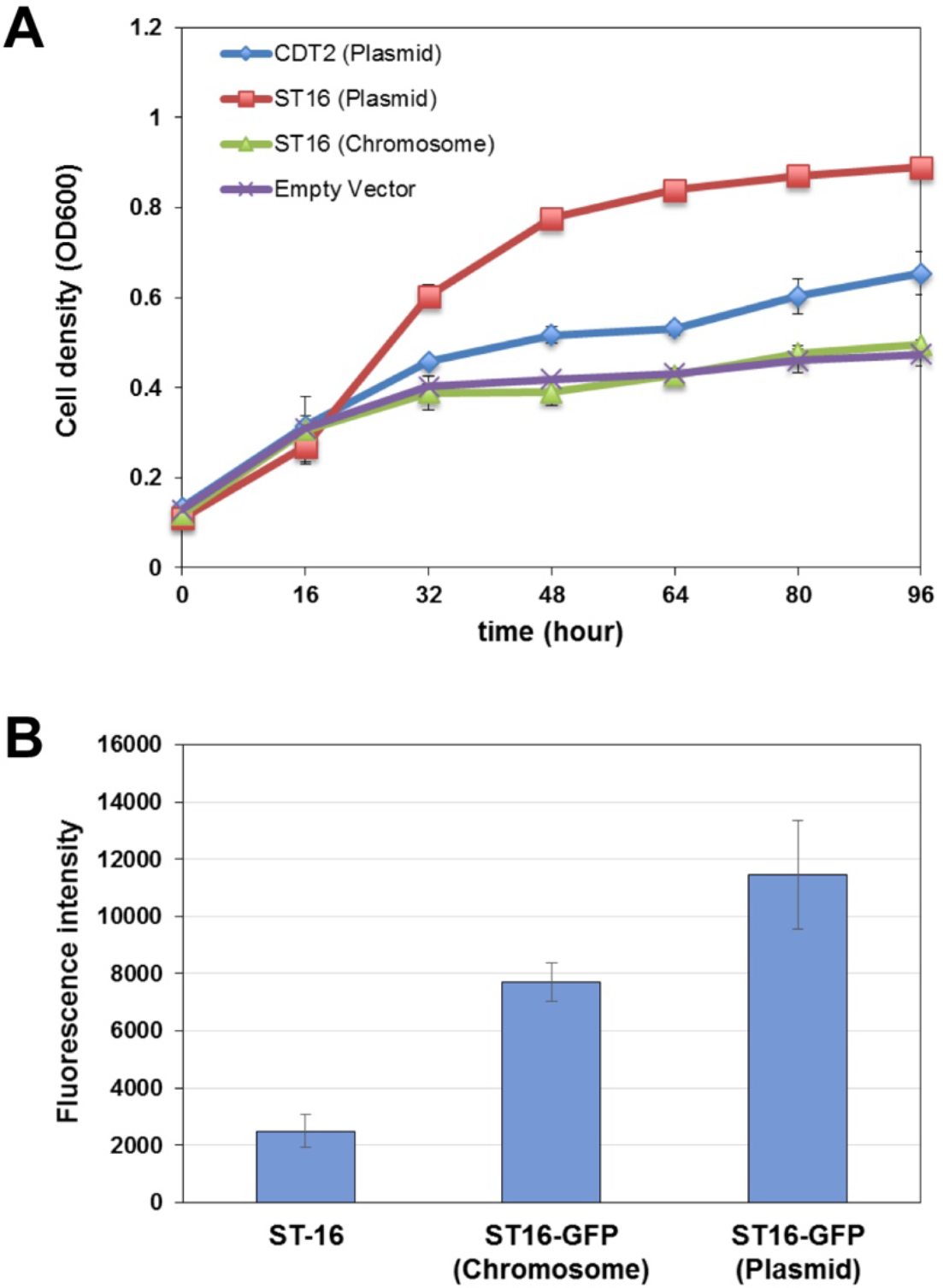
Chromosomal expression of transporter ST16 relative to plasmid-based expression slows aerobic growth. **A**). Aerobic growth profiles of yeast SR8A strains with plasmid expressing CDT-2, ST16 or empty control were compared with ST16 transporter expressed from a chromosomal copy at the *LEU2* locus. Error bars represent standard deviations of biological triplicates. **B**). GFP fluorescence emission spectroscopy analysis of ST16-GFP fusion expression. ST16, cells lacking the GFP fusion to establish baseline fluorescence.

To further test whether transport is limiting in XD-utilizing strains, we compared the aerobic growth profile of the SR8 strain with a plasmid expressing CDT-2, GH43-2 and GH43-7 and the SR8A strain, which has GH43-2 and GH43-7 expressed from chromosomal copies, with a plasmid only expressing the CDT-2 transporter. We reasoned that incorporation of the genes for GH43-2 and GH43-7 into the chromosome would drop the protein expression level compared with overexpression using a plasmid, because of the much higher copy number of plasmids [20, 24]. As expected for XD transport being the limiting factor for XD utilization, expressing GH43-2 and GH43-7 from the chromosome did not affect aerobic growth on xylodextrin (S3 Figure). These results further supported that the XD transporter is the rate-limiting step of the XD utilization pathway.

Next, we attempt to develop a CRISPR-Cas9 mediated directed evolution selection experiment [20] using growth in liquid media followed by screening based on colony size. We first used error prone PCR to generate a mutant library of ST16 of 2x10^3^ variants and transformed the library into the SR8A strain. Pooled strains from YPAD+G418 plates were grown aerobically in liquid synthetic media containing XD for 2 days and plated on plates with synthetic media containing XD to enrich for functional ST16 mutants. Surprisingly, after 4 days of aerobic growth on plates, all of the colonies had similar sizes, and we were not able to identify any individual colony that demonstrated faster aerobic growth. Furthermore, we conducted a serial dilution of xylodextrin concentration in making the synthetic plates, and found that the major roadblock with the selection strategy is the background growth of the strains on contaminating xylose and amino acids (S4 Figure), resulting in difficulty identifying functional ST16 mutants. Thus, the yeast strain was able to grow on alternative carbon sources rather than XD, circumventing the selective pressure of XD-dependent growth.

## Discussion

CDT-1 and CDT-2 are two important cellodextrin transporters used by *N. crassa* in cellulose degradation [25], of which only CDT-2 is also involved in hemicellulose utilization [7, 26]. In previous results, the discovery of the xylodextrin consumption pathway consisting of transporter CDT-2 along with two β-xylosidases GH43-2 and GH43-7 provided new modes for utilization of xylose derived from hemicellulose [7]. However, the XD pathway as isolated from *N. crassa* requires significant improvement for its future use industrially. In our study, we confirmed that the XD transporter is the rate-limiting step of the XD utilization pathway (Figure 6A and S3 Figure). Through characterization of a library of CDT-1 and CDT-2 transporter orthologues, we identified three potential transporters from other fungi that could be used as targets for exploring faster xylodextrin utilization (S2 Table). Of these, the transporter ST16 isolated from *Trichoderma virens* has advantages over CDT-2, including the fact that it is a XD-specific transporter with good membrane localization, and enables faster aerobic growth. Furthermore, the xylobiose transport activity of ST16 is not inhibited by the presence of cellobiose.

Given the superior properties of the ST16 transporter, we attempted to develop a CRISPR-Cas9 mediated directed evolution experiment, using ST16 as the starting point to screen mutants that could further improve xylodextrin utilization efficiency. We used liquid medium containing XD for selection, and subsequent screening for larger colonies on XD-containing plates. Through mutagenesis and chromosomal integration, we screened 2x10^3^ ST16 mutants. However, due to the background growth of these strains on xylose and amino acids, it was hard for us to select large colonies, resulting in difficulty in identifying more functional ST16 mutants. To overcome remaining bottlenecks of the current selection strategy, it will be necessary to remove the contaminating xylose from XD preparations, and minimize the amino acid concentrations in the media. In addition, further experiments need to be done for increasing the expression level of chromosomally integrated ST16, as we found that using the *PGK1* promoter was not sufficient to drive growth above background levels on XD-containing plates. Although not ideal, it may be necessary to first carry out directed evolution using ST16 expression from a plasmid, to increase ST16 expression levels. Other strategies might also be explored for directed evolution of XD transporters, including starting with ST2 or ST15. Alternatively, it might prove useful to perform a directed evolution experiment on a cellodextrin transporter like CDT-2 to improve cellobiose uptake, then test whether isolated mutations will also facilitate faster XD consumption.

Although both ST2 and ST16 did not improve anaerobic utilization of XD compared with CDT-2, the presence of xylose or other simple sugars could be used to initiate the consumption of XD, according to our previous results [7]. These anaerobic fermentation experiments suggest that metabolic sensing might require additional tuning for optimal xylodextrin fermentation. Based on the rapid turn-off and turn-on of XD consumption in anaerobic conditions [7], we suspect that the XD transporters are internalized in the absence of hexose or xylose sugars [27]. Currently, how CDT-2 and ST16 are regulated in xylodextrin-only media remains unknown. Other systems-level experiments such as transcription and ribosome profiling could also be used to better understand the mechanism by which the yeast strain senses xylodextrins under anaerobic conditions. Finally, the cofactor imbalance problem of the XR/XDH pathway may lead to the accumulation of xylose in the culture supernatant [28], indicating that the metabolic sensing and xylose assimilation pathway might require additional tuning for optimal xylodextrin fermentation.

## Materials and methods

### Strains and plasmids

The *N. crassa* and the codon-optimized versions of all ST transporters were cloned into the pRS316 plasmid (*CEN URA3*)—under the control of the *S. cerevisiae TDH3* promoter and the *CYC1* terminator—using the In-Fusion HD Cloning Kit (Clontech Laboratories, Inc., Mountain View, CA). The cleavage site for the HRV 3C protease followed by eGFP was fused to the C-terminus. *cdt-1 and cdt-2* were PCR-amplified from cDNA synthesized from mRNA isolated from *N. crassa* (FGSC 2489) grown on minimal media plus Avicel (microcrystalline cellulose) as the sole carbon source [25]. Atum (formerly DNA 2.0) performed gene synthesis and codon-optimization for all ST transporters. *S. cerevisiae* strain D452-2 [29], was used for XD and cellobiose transport studies. Codon-optimized *gh1-1* was cloned into pRS315 plasmid (*CEN LEU*) and was expressed under the control of the *PGK1* promoter. Details of the codon optimization process for *gh1-1* are described in [30].

*S. cerevisiae* strain SR8 [23] was used for strain engineering. We used CRISPR-Cas9 genome editing to insert GH43-2 and GH43-7 into the *TRP1* and *LYP1* loci, respectively, and the engineered SR8 strain was named SR8A. GH43-2 was under the control of *TDH3* promoter and the *ADH1* terminator, while GH43-7 was drove by *CCW12* promoter and the *CYC1* terminator.

### Growth conditions

For yeast growth experiments, optimized Minimal Media (oMM) lacking uracil or uracil plus leucine [30] was used. oMM contained 10 g/L (NH4)2SO_4_, 1 g/L MgSO_4_ · 7H_2_O, 6 g/L KH_2_PO_4_, 100 mg/L adenine hemisulfate, 1.7 g/L YNB (Sigma-Aldrich, Y1251), 2x recommended CSM-Ura dropout mix (MP Biomedicals, Santa Ana, CA), 10 mg/L of inositol, 100 mg/L glutamic acid, 20 mg/L lysine, 375 mg/L serine, 100 mM morpholineethanesulfonic acid (MES), pH 5.7. Cellobiose or xylodextrin was added to this stock recipe depending on the experiment.

For directed evolution experiments, strains were grown at 30 °C in either rich (yeast extract-peptone [YP]) or synthetic (S) medium containing 1% xylodextrin with appropriate nutrient supplements to support growth and with certain nutrients omitted to maintain selection for plasmids. Xylodextrin was purchased from Cascade Analytical Reagents and Biochemicals. The composition of the XD samples was analyzed by Dionex ICS-3000 HPLC (ThermoFisher) as described below. Synthetic media contained 2 g/L yeast nitrogen base without amino acids and ammonium sulfate, 5 g/L ammonium sulfate, 1 g/L CSM.

### Aerobic growth assays

All growth assays were performed using the Bioscreen C (Oy Growth Curves Ab Ltd., Helsinki, Finland) with biological triplicates or quadruplicates. Single colonies of *S. cerevisiae* strains transformed with pRS316 containing the MFS transporter of interest were grown in oMM-Ura plus 2% glucose to late-exponential phase at 30 °C in 24-well plates. Cultures were pelleted at 4000 rpm, the spent media supernatant was discarded and cells were resuspended in H_2_O. The OD was measured to calculate the inoculum volume needed for 200 μL cultures at an initial OD ≈ 0.1 or 0.2 in Bioscreen plates in oMM media. The OD at 600 nm was measured in 30 min intervals for 48-96 h at 30 °C.

### Epi-fluorescence Microscopy

D452-2 cells expressing the transporters were grown in oMM-Ura plus 2% glucose media to mid-exponential phase at 30 °C. The cultures were centrifuged, spotted onto glass slides and examined on a Leica DM 5000B Epi-fluorescence microscope at 100x DIC (Leica, Germany). Transporters were visualized using the L5 filter cube; images were captured using the Leica DFC 490 camera and analyzed with the accompanying microscope software.

### Yeast-cell based sugar uptake assay

Yeast strain D452-2 cells transformed with pRS316-*CDT1, -CDT2* or pRS316-*ST* were grown to mid-exponential phase. Cells were harvested and washed twice with Transport Assay Buffer (5 mM MES, 100 mM NaCl, pH 6.0) and resuspended to a final OD_600_ of 40. 500 μL of cell resuspension was quickly mixed with an equal volume of Transport Assay Buffer containing 400 μM of the respective sugar (final sugar concentration was approximately 200 μM). For the initial time point (t = 0 sec), an aliquot was removed and centrifuged for 1 min at 4 °C at high speed to pellet the cells and the supernatant was removed. The remaining cell resuspension was incubated at 30 °C for 30 min with constant shaking. After incubation, samples were centrifuged for 5 min at 4 °C at 14000 rpm and supernatant was removed. For analysis, 400 μL of supernatant were mixed with 100 μL of 0.5 M NaOH, and sugar concentrations remaining in the supernatant were measured by Dionex ICS-3000 HPLC as described below.

### HPLC analysis

HPLC analysis was performed on Dionex ICS-3000 HPLC using a CarboPac PA200 analytical column (150 x 3 mm) and a CarboPac PA200 guard column (3 x 30 mm) at room temperature. 25 μL of sample was injected and run at 0.4 mL/min with a mobile phase using 0.1 M NaOH. Acetate gradients were used to resolve xylodextrin samples. For xylobiose and xylotriose, the acetate gradients were as follows: 0 mM for 1 min, increasing in 8 min to 80 mM, then to 300 mM in 1 min, keeping at this concentration for 2 min, followed by equilibration back to 0 mM for 3 min.

### CRISPR-Cas9 genome editing

CRISPR-Cas9 genome editing was performed according to published precedures [20] with slight modifications. In brief, Cas9 transformation mix consisted of 90 μL yeast competent cells (OD600 ≈ 1.0), 10.0 μL of 10 mg/mL ssDNA (D9156, Sigma, St. Louis, MO), 1.0 μg pCAS plasmid, 5.0 μg of linear repair DNA and 900 μL Polyethyleneglycol 2000 (295906, Sigma), 0.1 M Lithium acetate, 0.05 M Tris–HCl and 5.0 mM EDTA. The mix reagents were incubated 30 min at 30 °C, and then subjected to heat shock at 42 °C for 17 min. Following heat shock, cells were centrifuged at 5000 rpm for 2 min. The pellet was resuspended in 250 μL YPD and recovered for 2 hour at 30 °C, then the entire contents were plated onto YPAD+G418 plates (20 g/L Peptone, 10 g/L Yeast Extract, 20 g/L Agar, 0.15 g/L Adenine hemisulfate, 20 g/L Glucose and 200 mg/L G418). Cells were grown 24 hours at 37 °C, then moved to 30 °C to complete growth. Colonies from the YPAD+G418 plates were picked and grown overnight in 1.0 mL of liquid YPAD medium. Genomic DNA was extracted from these cultures using the MasterPure Yeast DNA Extraction Kit (MPY80200, Epicentre). PCR confirmation of the integration allele was performed, and PCR products were submitted for Sanger sequencing at the UC Berkeley, Sequencing Facility (Berkeley, CA) to confirm the integration sequence.

### Protein expression levels using fluorescence emission spectroscopy

For comparison of ST16 expression from plasmid-encoded and chromosomally-integrated genes, cells were grown in 10 mL of SC-Ura media (Sunrise Science Products) with 1% glucose under aerobic conditions at 30 °C overnight. Glucose was used rather than XD, due to the very slow growth of cells in XD-containing media. Cells were normalized to a total OD of 20.8 in 200 μL and measured for GFP fluorescence signal using the Synergy Mx plate reader (BioTek) with the following filter set: excitation 485/20, emission 528/20. Cells expressing ST16 without the GFP fusion were used as a control to establish the baseline fluorescence signal from the yeast cells.

For comparisons of transporter expression from plasmids (S5 Figure), 50 mL cultures were grown at 30 °C until they reached an OD_600_ of 3, at which point they were harvested by centrifugation, resuspended in ~4-5 mL of media and aliquoted into microcentrifuge tubes to yield a total OD_600_ of 30. Samples were spun down at 14,000 rpm for 1 min and the supernatant was aspirated. Cell pellets were quickly flash frozen in liquid N_2_. Frozen cell pellets were thawed on ice and 400 μL of Buffer A (25 mM Hepes, 150 mM NaCl, 10 mM cellobiose, 5% glycerol, 1 mM EDTA, 0.2X HALT protease inhibitor cocktail (ThermoFisher, Waltham, MA), pH 7.5) were added for resuspension. Cells were lysed with zirconia/silica beads in a Mini-Beadbeater-96 (Biospec Products, Bartlesville, OK). Cell debris was pelleted at 10,000x*g* for 10 min at 4 °C, lysates were diluted three-fold with Buffer A, and their GFP fluorescence was measured using a Horiba Jobin Yvon Fluorolog fluorimeter (Horiba Scientific, Edison, NJ). The λ_EX_ was 485 nm, and the emission wavelength was recorded from 495-570 nm, with both excitation and emission slit widths set to 3 nm. A fluorescence calibration curve was prepared with eGFP purified from *E. coli* (>95% purity). The settings and the eGFP protein concentration range were chosen to yield a linear correlation between the fluorescence intensity at the maximum λ_EM_ (510 nm) and the protein concentration of the standard. The maximum fluorescence intensity of the samples fell within this range. Target protein concentrations represent the mean from 3 biological replicates. Total protein concentration of the lysate was determined using the Pierce BCA Protein Assay Kit (ThermoFisher).

## Acknowledgements

We thank David Nunn at British Petroleum for the library of CDT-1 and CDT-2 orthologues. This work was supported by the Energy Biosciences Institute. C.Z. was supported by the China Scholarship Council.

